# Prevalence of Stunting and Associated Factors among public Primary School pupils of Bahir Dar city, Ethiopia: School-Based Cross-Sectional Study

**DOI:** 10.1101/756239

**Authors:** Getasew Mulat Bantie, Kidist Hailu Akenew, Mahlet Tilahun Belete, Eyerusalem Teshome Tena, Genet Gebreselasie Gebretsadik, Aynalem Nebebe Tsegaw, Tigist Birru Woldemariam, Ashenafi Abate Woya, Amare Alemu Melese

## Abstract

**Background:** Stunting is a well-established pupils-health indicator of chronic malnutrition which reliably gives a picture of the past nutritional history and the prevailing environmental and socioeconomic circumstances.

**Objective:** The prevalence of stunting and associated factors among public primary school pupils of Bahir Dar city.

**Method:** A cross-sectional study was carried out from March to June 2019. Data were coded and entered into Epi-Data and exported to SPSS. Then, Anthropometric data were converted into height for age Z-scores to determine the pupils stunting outcomes using WHO Anthro-Plus software. Then, the final analysis was done by SPSS version 20 software. Anthropometric measurements determined the proportion of stunting (z-score of height for age less than minus two standard deviations from WHO Anthro-plus software output). A simple logistic regression model was fitted to identify factors associated between the independent variables and the dependent variable at a 95% confidence level and p-value <0.05.

**Results:** 370 primary school pupils were included in the study with the mean age of 121.84(± 26.67) months. About 51.6% of the pupils were females. The total prevalence of stunting was 15.13% (95%CI; 11%, 19%). The burden of stunting was higher in the age group of greater than 132 months. Pupil’s age ≥132 months (AOR=15.6; 95%CI; 3.31, 73.45; p-value<0.001) and male pupils (AOR=7.07; 95%CI: 2.51, 19.89; p-value<0.0002) were significantly associated with stunting.

**Conclusion:** The prevalence of stunting was relatively lower than the regional estimated stunting level. However, this result is also very significant figure to get critical attention. Pupil’s age ≥ 132 months and male sex were significantly associated with stunting.

## INTRODUCTION

School pupils pass through great physical and mental changes, which affect both their growth and school performance(1) and nutrition is one of the many key factors affecting mental development of children (2).

Undernutrition is a condition that results from eating a diet which has either inadequate nutrient to the extent that causes health problems(3). Under nutrition is widespread among school children particularly in low income countries particularly in Africa and their nutritional status often worsens during their school years. It greatly affects both the cognitive and physical development of school age children(4). It is known that it may increase susceptibility to infection(5) undernutrition among children has lifelong impacts; its effect could also transfer from the current generation to future.

Under nutrition related health challenges in school-age children are among the most frequent reasons of low school enrolment, high absenteeism, early dropout, poor academic performance, delayed cognitive development, short stature, reduced work capacity, and poor reproductive performance(4) According to some studies done in rural Ethiopia, the extent of stunting in school-age children ranges from 26.5% to 42.7%(6). Among the urban dwellers of this group, the range of stunting was 5.4% to 29.2(7).

Different studies have been done on factors determining undernutrition, but still, there are inconsistencies across the studies especially primary school children.

Therefore, the main aim of this study is to determine prevalence of stunting and associated factors among public primary school children of Bahir Dar city. The results of this study may reveal the existing prevalence and determinants of stunting among public primary school children of Bahir Dar city.

## METHODS AND MATERIALS

### Study design, setting and period

A school-based cross-sectional study was conducted in Bahir Dar city public primary school pupils from March 10 to June 10, 2019. Bahir Dar city is located 565 km away from Addis Ababa, the capital city of Ethiopia. The city is administratively had six sub-cities. There are two governmental and two private hospitals, eight governmental health centers, thirty-six private clinics and one governmental University, four private and two governmental colleges are found in the city. The total population of the city is estimated to be 249,851 (124,426 females and 125,425 males), according to the population and household survey. According to Bahir Dar city education department of 2018/19 report, there are 19 public primary schools (1 up to 8 graded).

### Population, sample and sampling procedure

The source population for this study consisted of all pupils attending primary schools of Bahir Dar city. The study population for this study was pupils attending the selected primary schools of the city were included in the study. Those with drop out or absent during the data collection period were excluded. The sample size was determined using a single population proportion formula by considering; 95% confidence level, 5% margin of error, stunting level of 18.3% in Bahir Dar city(8). Taking the design effect 1.5 and 10% non-response rate and considering the correction formula (N<10,000), the final sample size was 375. Then, multistage sampling was used to select eligible pupils. Five primary schools were recruited for the study from 19 total primary schools (1 up to 8 graded) by lottery method. From the five selected schools, primary cycle (1 up to 4 graded) pupils were recruited for the study. Then, systematic sampling technique with proportional allocation to size was applied from the registration list to select 375 samples. **(Figure 1)**

**Figure 1:**
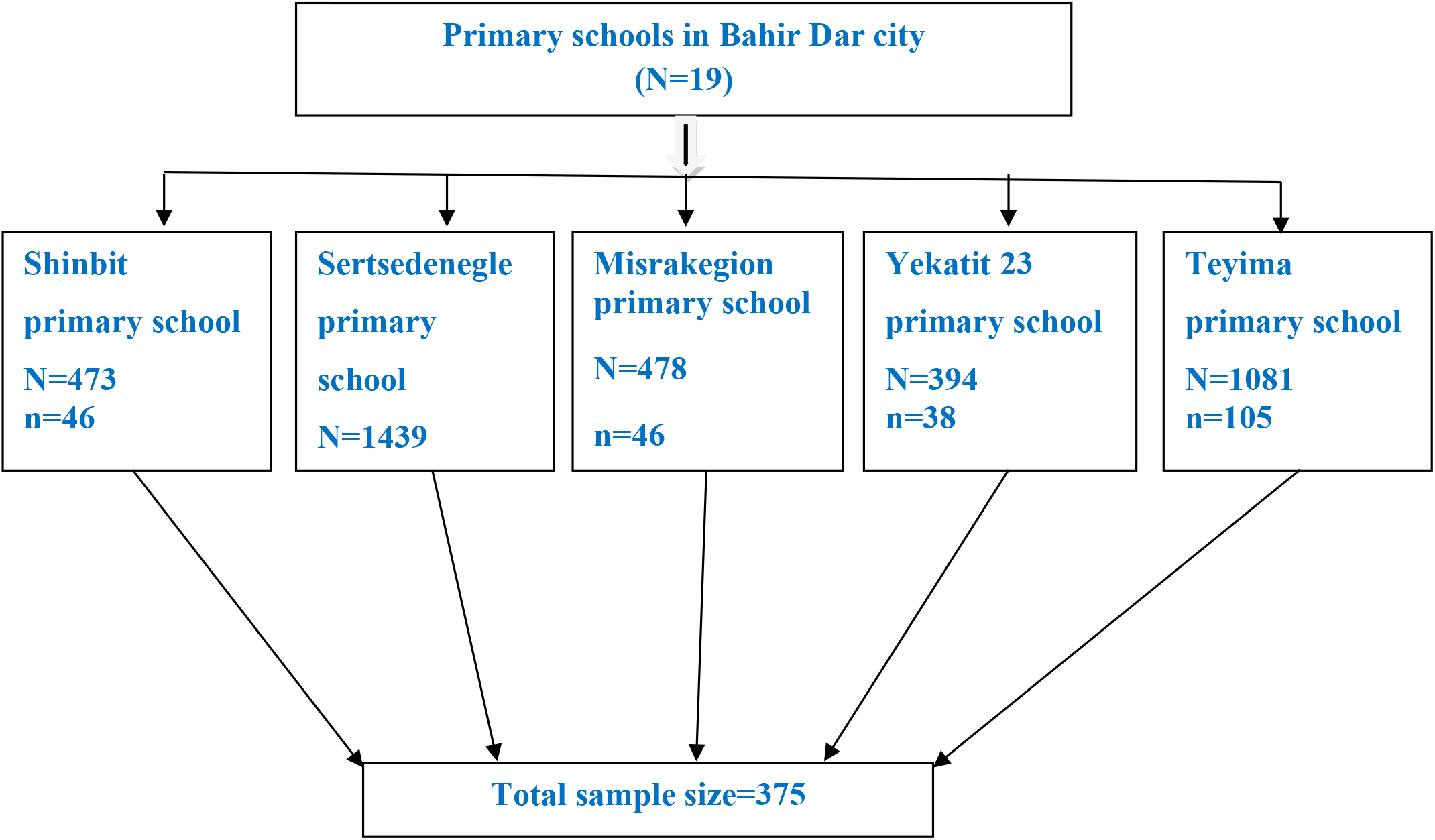
Schematic presentation of the sampling procedure in Bahir-Dar city, 2019.

### Data collection procedures and measurement of variables

Data collection tools initially prepared in English then translated to local Amharic language. Six BSC nurses (four for data collection and two for supervision) were recruited. Two days training was given for data collectors and supervisors by the principal investigator about the objective of the study. Socio-demographic characteristic, health care characteristic, maternal awareness on dietary diversification, and household food security status-related data were collected using a pretested structured interviewer-administered questionnaire. Anthropometric data such as weight, height, and body mass measurements were taken from each study participants. After anthropometric measurements were taken, study participants’ parents were interviewed with face to face based on the objective of the study. Participants were classified as stunted if they had z-score of height for age less than minus two standard deviations from WHO Anthro-plus software output.

### Data Processing and analysis

Data were coded and entered into Epi-Data and exported to SPSS version 20 software and then, to WHO Anthro-Plus software. Again back to SPSS for analysis. Descriptive statistics were used to summarize characteristics of study participants and presented using text and table. Bivariate and multivariate logistic regression analysis was employed to assess the association between the exploratory variables and stunting. The strength of the association was measured using the adjusted odds ratio (AOR) and 95% confidence interval (CI). A p-value < 0.05 was considered as a statistically significant predictor of stunting.

## RESULT

### Socio-demographic characteristics of the participants

A total of 370 primary school pupils were recruited for the study with a response rate of 98.6%. The mean age of the respondents was 121.84(+ 26.67) months. About 51.6% of the stunted pupils were females. About 35% of households with more than ten family size got their child stunted. 20% of the merchant parents had a stunted child. About 15% of the mothers were not able to read and write and 20% had family monthly income of fewer than 500 birrs. Similarly, more than 15% of the child was underweight (15.5%) and wasted (17.9%), respectively. **(Table 1)**

**Table 1:** Socio-demographic related characteristics of the participants, Bahir Dar city, North West Ethiopia, 2019

| Variable | Category | Stunted | Not-stunted |
| --- | --- | --- | --- |
|  |  | N(%) | N(%) |
| Age of the pupil<br>(in months) | 72-96 | 90(92.8) | 7(7.2) |
|  | 108-120 | 116(89.2) | 14(10.8) |
| | ≥ 132 | 108(75.5) | 35(24.5) |
| Sex of the pupil | Female | 22(11.5) | 169(88.5) |
|  | Male | 34(19.0) | 145(81.0) |
| Religion | Orthodox | 49(14.7) | 283(85.3) |
|  | Islam | 7(22.6) | 24(77.4) |
|  | Catholic | 0(0) | 2(100) |
|  | Protestant | 0 | 5(100) |
| Family size of the household | 1-5 | 27(13.6) | 172(86.4) |
|  | 6-10 | 22(14.6) | 129(85.4) |
|  | 11-15 | 7(35) | 13(65) |
| Mother's occupation | Merchant | 98(20) | 36(80) |
|  | Government employee | 5(11.4) | 39(88.6) |
|  | Self-employed | 26(18.3) | 116(81.7) |
|  | Housewife | 15(11.1) | 120(88.9) |
| Mother's educational status | Can't read and write | 18(15.5) | 98(84.5) |
|  | Can read and write | 23(17.8) | 106(82.2) |
|  | Primary school complete | 7(11.9) | 52(88.1) |
|  | Secondary school and above | 7(11.3) | 55(88.7) |
| Monthly income( in Birrs) | <500 | 12(20.3) | 47(79.7) |
|  | 500-1000 | 13(14.1) | 79(85.9) |
|  | 1000-1500 | 8(13.6) | 51(86.4) |
|  | > 1500 | 23(14.4) | 137(85.6) |
| Body mass index | Under weight | 42(15.3) | 232(84.7) |
|  | Normal | 9(15.2) | 50(84.8) |
|  | Over weight | 1(50) | 1(50) |
| Middle upper arm circumference | Wasting | 27(17.9) | 124(82.1) |
|  | Normal | 29(13.2) | 190(86.8) |

### Household food security, dietary diversification, and health care

22.2% of the parents who were worried for fear of not having enough food in the household had their child stunted. Similarly, those parents who shared foods from relatives when a shortage of food happens were their child stunted. Family members who didn’t get enough food were 19% their child stunted. 15% of stunted children seen in households faced a shortage of food. The main reason, (24.1%) for this shortage was due to low family income were stunted.

24.5% of the stunted children mothers’ did not participate in their child feeding. The families who didn’t hear about the variety of food were 21.1% their child stunted. 17.6% of stunting was observed in the family members who had a chronic disease. Similarly, 16.7% of currently ill study children were stunted **(Table 2)**

**Table 2:** Household food security, dietary diversification, and healthcare-related characteristics of the participants, Bahir Dar city, North West Ethiopia, 2019

| Variable | Category | Stunted<br>N(%) | Not-stunted<br>N(%) |
| --- | --- | --- | --- |
| Did you worry for fear of not having enough* household food | Yes | 35(22.2) | 123(77.8) |
|  | No | 21(9.9) | 191(90.1) |
| If the family get a shortage of food what is your measurement | Share with relatives | 12(22.2) | 42(77.8) |
|  | Debt | 14(13.2) | 92(86.8) |
|  | Work hard | 20(16.1) | 104(83.9) |
|  | Do nothing | 4(9.3) | 39(90.7) |
|  | Asking for donation | 6(14) | 37(86) |
| In your judgment is there anyone who didn't get enough food* in your family | Yes | 23(19) | 98(81) |
|  | No | 33(13.7) | 216(86.3) |
| Do you face shortage of food | Yes | 22(15) | 125(85) |
|  | No | 34(15.2) | 189(84.8) |
| Reason for food shortage | Divorce | 4(10.9) | 33(89.1) |
|  | Fool about | 2(8) | 23(92) |
|  | In come shortage | 14(24.1) | 44(75.9) |
|  | Lack of peace in the county | 0(0) | 13(100) |
|  | Economical in stability | 2(14.3) | 12(85.7) |
| The mother participate in child feeding | Yes | 44(13.7) | 277(86.3) |
|  | No | 12(24.5) | 37(75.5) |
| Do you hear about variety of food | Yes | 52(14.8) | 299(85.2) |
|  | No | 4(21.1) | 15(78.9) |
| Source of information | Health care provider | 22(13.8) | 138(86.3) |
|  | Family members | 5(8.1) | 57(91.9) |
|  | Television | 17(23.0) | 57(77.0) |
|  | Radio | 3(21.4) | 11(78.6) |
| Variety of food is good for your children | Yes | 55(15.1) | 309(84.9) |
|  | No | 1(16.7) | 5(83.3) |
| Where will you go when your child gets sick | Health institution | 39(14.8) | 225(85.2) |
|  | Traditional medicine | 4(10.5) | 34(89.5) |
|  | Holly water | 13(19.7) | 55(80.3) |
| Family member who had any chronic disease** | Yes | 6(17.6) | 30(83.3) |
|  | No | 50(15.0) | 284(85.0) |
| Current health status of this child | Healthy | 4(15.0) | 20(85.0) |
|  | Ill | 52(16.7) | 294(83.3) |

Enough food\*(eating food at least three times per day); chronic disease\*\* (diabetes mellitus, hypertension and/or asthma)

### The prevalence of stunting among primary school pupils

This study revealed that 15.13% of the pupils of public primary school of the city were stunted. (Figure 2)

**Figure 2:**
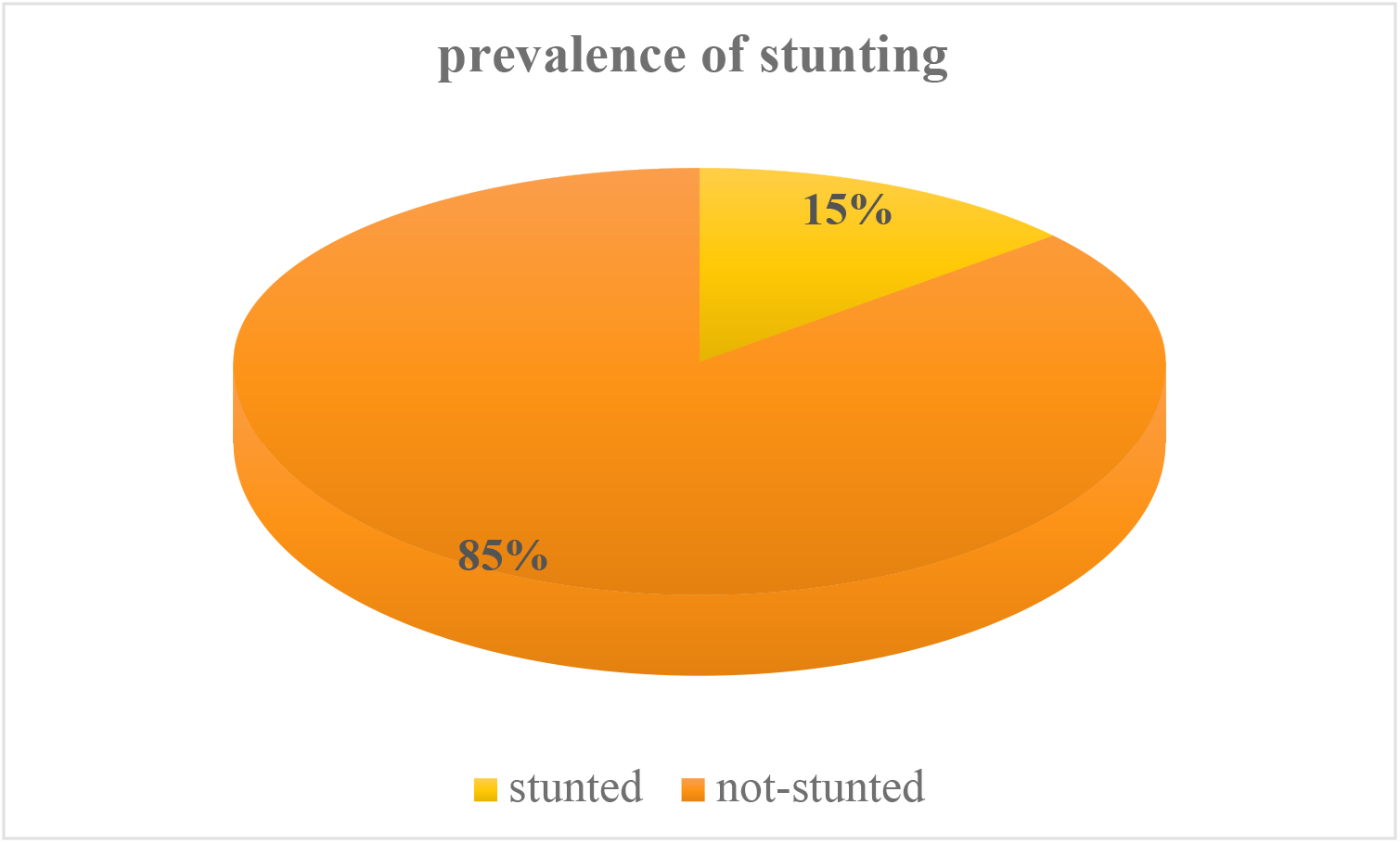
the prevalence of stunting among primary school pupils of Bahir-Dar city, 2019.

### Factors associated with stunting among primary school children

Multivariable logistic regression analysis was conducted to identify independent predictors of stunting among public primary school pupils. On multivariable logistic regression model, age and sex of the pupil were significantly associated with stunting at a p-value of 0.05.

Accordingly, those primary school children in the age group of > 132 months were about 15 times, (AOR=15.6: 95%CI: 3.31, 73.45) more likely to be stunted compared to 72-98 months’ of age group. The odds of stunting among male pupils were about 7 times (AOR=7.07; 95%CI: 2.51, 19.89) higher compared to those female pupils. **(Table 3)**

**Table 3:** Factors associated with stunting among primary school children in Bahir Dar city, North West Ethiopia, 2109

| Variable | Stunting |  | COR (95%CI) | AOR (95%CI) | P-value |
| --- | --- | --- | --- | --- | --- |
|  | Yes | No |  |  |  |
| Age of the child (in months) |  |  |  |  |  |
| 72-96 | 90 | 7 | 1.00 | 1.00 |  |
| 108-120 | 116 | 14 | 1.55(0.6,4.0) | 4.44(0.88,22.34) | 0.070 |
| ≥ 132 | 108 | 35 | 4.17(1.76,9.83) | 15.6(3.31,73.45) | 0.001 |
| Family size |  |  |  |  |  |
| 1-5 | 27 | 172 | 1.00 | 1.00 |  |
| 6-10 | 22 | 129 | 1.08(0.59,1.99) | 1.83(0.72,4.67) | 0.199 |
| 11-15 | 7 | 13 | 3.43(1.26,9.36) | 2.95(0.43,19.81) | 0.266 |
| Worrying due to fear of not having enough household food |  |  |  |  |  |
| Yes | 35 | 123 | 2.58(1.44, 4.65) |  |  |
| No | 21 | 191 | 1.00 | 1.00 |  |
| Frequency of worrying due to fear of not having enough household food |  |  |  |  |  |
| Always | 10 | 27 | 1.00 | 1.00 |  |
| Usually | 7 | 57 | 0.33(0.11,0.96) | 0.45(0.13,1.25) | 0.116 |
| Sometimes | 35 | 123 | 1.24(0.49,3.11) | 1.75(0.64,4.79) | 0.274 |
| Current health status of the pupils |  |  |  |  |  |
| Healthy | 4 | 20 | 1.00 | 1.00 |  |
| Ill | 52 | 294 | 1.13(0.37,0.44) | 1.83(0.41,8.08) | 0.426 |
| Sex of the pupil |  |  |  |  |  |
| Female | 22 | 169 | 1.00 | 1.00 |  |
| Male | 34 | 145 | 1.80(1.00,3.21) | 7.07(2.51,19.89) | 0.0002 |

## DISCUSSION

The present study revealed that the prevalence of stunting among primary school pupils in Bahir Dar City Administration was 15.13% (95%CI; 11, 19). This finding was consistent with studies conducted at Rabwah, Pakistan, 11.5%(9) and Baghdad, Iraq, 18.7%(10), Southern Pakistan,16.5%(11), in Abeokuta, Southwest Nigeria,17.4%(12)

But the present result showed a lower prevalence than a study conducted at Jimma Zone, southwest Ethiopia,24.1% (13), and Urban Communities in Obafemi Owode, Southwestern Nigeria, 33.8%(14), Gondar town, Ethiopia, 46.1% (15), Humbo district, Southern Ethiopia, 57%(16), west Bengal,23.0%(17), Western Kenya,36% (18), Mecha District, Amhara Regional State, 37.9%(19), southern Ethiopia,28%(20),southern Angola, 41.5%(5), in Cambodian, 40.0%(2), in Arba Minch Health and Demographic Surveillance Site, Southern Ethiopia,41.9%(4)

On the contrary, the current study finding was higher than the study finding from Northcentral Nigeria 10.5%,(21), Tehran, Iran,3.7%(22), in China, 1.0-1.9%(23), Bogota, Colombia,19.8%(24), northern region of Sudan,7.1%(25), Argo city, Northern Sudan, 3.8%(1), in Lahore, Pakistan, 8%(26), in Jiangsu Province, China, 4.7%(27), in Beijing, China, 2.6%(28). The possible explanation might be the difference in socio-demographic characteristics like study setting and lifestyle and health information dissemination.

The current study also identified the risk factors associated with stunting among public primary school pupils in Bahir Dar city.

Pupils in the age group of 132 months and above were sixteen times more likely stunted than the age group of 72-96 months. This finding was supported by a study done in Southern highlands of Tanzania (29), Southwestern Nigeria(14), Baghdad, Iraq(10), Humbo district, Southern Ethiopia(16), Gondar town, Ethiopia(15), Southern Pakistan(11), Mecha District, Amhara Regional State(19), Lahore, Pakistan(26), Southern Angola(5),southwest Nigeria(12), and Arba Minch, Ethiopia(4). This might be due to the fact older children may suffer from the presence of younger siblings.

Male primary school pupils were sevenfold at higher risk for stunting than females; which was supported by a study finding from Southern highlands of Tanzania(29), and Arba Minch, Ethiopia (4) and EDHS 2016(30). In contrast, the study finding from Beijing-China(28) and Southern Pakistan,(11) revealed that females were more likely at risk of being stunted than males. This might be because males’ growth and development is more influenced by environmental and nutritional stress than females and thus, making males more likely to be affected by stunting (4).

### Limitation of the study

There might be a potential bias among respondents answering questions relating to events happening in the past such as food availability. Measurement bias might also occur during data collection.

### Conclusion

The prevalence of stunting was relatively lower than the regional estimated stunting level. However, this result was a very significant figure to get critical attention. Pupil’s age ≥ 132 months and male sex were significantly associated with stunting.

## Acronyms/Abbreviations

AOR: Adjusted Odds Ratio
BMI: Body Max Index
CI: Confidential Interval
EDHS: Ethiopian Demographic and Health Survey
SD: Standard Deviation
WHO: World Health Organization

## Ethical considerations

Ethical clearance was obtained from GAMBY Medical and Business College, Research and Publication Office with the reference number of GC-221/2011. The support letter was obtained from Bahir Dar city health and education department before the beginning of data collection. The directors of each primary school was informed about the purpose of the study that it will contribute for the health promotion and intervention of the primary school children of the study area and in the nation at large. Before collecting the data, parents of the students were informed and written consent was obtained from them and finally the parents were interviewed. Names of the students and/ or parents did not use to ensure anonymity and confidentiality.

## Availability of data and materials

The data can be accessed from the corresponding author

## Consent to publish

Written consent was obtained that the interview will be included in publications

## Competing interests

Authors declare that there are no competing interests.

## Funding

No fund was obtained

## Authors’ Contributions

GMB, KHA, MTB, ETT, GGG, ANT, TBW AAW, and AAM involved in the design, data collection, statistical analysis and interpretation, and manuscript drafting and critical revision. All authors have an equal contribution.

## Acknowledgments

We would like to thank our data collectors and the supervisor for their invaluable effort; our deep gratitude also goes to our study participants who volunteered and took their time to give us all the relevant information for the study. Last but not least, we would like to thank all Bahir Dar city primary school directors for their cooperation and support during the data collection.

